# RNA velocity in single cells

**DOI:** 10.1101/206052

**Authors:** Gioele La Manno, Ruslan Soldatov, Hannah Hochgerner, Amit Zeisel, Viktor Petukhov, Maria E. Kastriti, Peter Lönnerberg, Alessandro Furlan, Jean Fan, Zehua Liu, David van Bruggen, Jimin Guo, Erik Sundström, Gonçalo Castelo-Branco, Igor Adameyko, Sten Linnarsson, Peter V. Kharchenko

**Author notes:** **Corresponding authors.** (S.L.) and (P.V.K.).

## Abstract

RNA abundance is a powerful indicator of the state of individual cells, but does not directly reveal dynamic processes such as cellular differentiation. Here we show that RNA velocity—the time derivative of RNA abundance—can be estimated by distinguishing unspliced and spliced mRNAs in standard single-cell RNA sequencing protocols. We show that RNA velocity is a vector that predicts the future state of individual cells on a timescale of hours. We validate the accuracy of RNA velocity in the neural crest lineage, demonstrate its use on multiple technical platforms, reconstruct the branching lineage tree of the mouse hippocampus, and measure RNA kinetics in human embryonic brain. We expect RNA velocity to greatly aid the analysis of developmental lineages and cellular dynamics, particularly in humans.

## Introduction

Single-cell RNA sequencing measures gene activity with high quantitative accuracy, sensitivity and throughput, and is increasingly used to discover the cellular composition of complex tissues (*1*). The distribution of cells in the expression space provides a partial reconstruction of the manifold on which physiological expression states of different cells can reside. In dynamic contexts, such as embryogenesis or regenerating tissues, it is the movement of cells along this expression manifold that is of primary interest. In other words, one is interested in reconstructing the temporal sequence of transcriptional changes leading up to any given cell fate. Single-cell RNA-seq, however, captures only a static snapshot of the cellular state at a point in time, posing a critical challenge for reconstruction of cellular dynamics.

Current experimental methods to trace cell fate and reconstruct cell lineages have limited power (*2*). Genetic methods to map the fate of defined cell types reveal only the end state, and not the intervening trajectory of differentiation. Time-lapse imaging can record lineages over short timescales (*3*), but does not reveal the molecular identity of intermediate states. In humans, genetic methods are not available, with the exception of naturally occurring somatic mutations, which however currently require expensive and low-throughput single-cell whole-genome sequencing (*4*, *5*).

Existing approaches to infer cell lineage and expression dynamics from single-cell RNA-seq snapshots have focused on the analysis of cell density, commonly modeling expression dynamics as progression of cells along an idealized central trajectory, such as a continuous curve or a tree. Under an assumption of a ergodic process, where the measurement snapshot has captured cells in all intermediate stages, central trajectories can be inferred by connecting all of the observed cell density in an optimal manner. An extensive set of different trajectory inference methods has been developed (*6*–*9*), and applied to modeling of cell differentiation or perturbation response (*6*, *8*, *10*–*12*). The resulting trajectories, however, do not provide any indication of the direction in which the cells progress, requiring explicit assumptions about the start and end points of the dynamic process. Similar limitations also hinder more advanced methods that model expression dynamics as being driven by a potential landscape, particularly in non-stationary contexts such as organism development, where boundary conditions can be difficult to define (*13*, *14*). Fundamentally, the problem is that observed static snapshots of cell distributions in expression space are compatible with multiple, conflicting, underlying dynamical processes (*13*).

During development of the mouse embryo, key differentiation events occur on timescales of hours to days. Human development is generally slower, while for example, zebrafish develops every organ in less than 24 hours. We noted that these timescales are comparable to the kinetics of the mRNA lifecycle – transcription, splicing, nuclear export, translation and degradation – with typical half-lifes on the order of ten hours, and that individual cells would simultaneously contain both newly synthesized mRNA, mature mRNA, and partially or completely degraded mRNA. We reasoned that by distinguishing mRNA molecules at different life-cycle stages, we might be able to observe both the present (mRNAs currently being translated into protein), past (mRNAs being degraded) and future (newly transcribed mRNAs) states of individual cells. Furthermore, mRNA molecules at different stages in the RNA lifecycle have distinguishable properties: newly transcribed mRNA has introns, spliced but not exported mRNAs reside in the nucleus, currently translated mRNA is bound to ribosomes, and degraded mRNA is partially fragmented. These features can all in principle be exploited to obtain a measure of transcriptional velocity, *i.e*. the rate of change of mRNA molecule abundances in the cell. Similar ideas have been previously leveraged to measure RNA kinetics in time-series microarray data, without the need for metabolic labelling (*15*). We reasoned that the manifolds observed in single-cell RNA-seq data could be analogous to an explicit time-series of differentiation.

We focused on splicing and degradation, which can be directly observed in the mRNA sequence itself, but found that the degradation signal was weak and confusing, possibly because RNA molecules can appear fragmented for technical reasons not having to do with active degradation (*e.g*. incomplete reverse transcription). Splicing, however, was readily detected by the presence of reads spanning exon/intron boundaries (indicating unspliced pre-mRNA) and reads spanning exon/exon boundaries (indicating spliced, mature mRNA). Reads mapping entirely inside exons could be assigned to either spliced or unspliced RNAs only approximatively, based on inter-exon distance, while reads mapping entirely inside introns had to be treated with caution to distinguish them from non-specific background and cryptic intron-located transcripts (see Methods).

All common single-cell RNA-seq protocols rely on oligo-dT primers to enrich for polyadenylated mRNA molecules. These should generally represent fully-spliced and polyadenylated mature mRNAs. Nevertheless, examining single-cell RNA-seq datasets based on the SMART-seq2, STRT/C1, in Drop, and 10X Chromium protocols (*16*–*19*), we found a substantial fraction (10-35%) of molecules containing unspliced intronic sequences (Fig. 1A). These included molecules originating entirely from the intronic regions, as well as molecules spanning the junction between exonic and intronic regions. Capture of precursor mRNAs has been previously observed in bulk RNA-seq data, suggesting that relative abundance of such molecules could be used to estimate relative rate of nascent transcription (*20*). The substantial number of spanning and intronic molecules, correlation of their abundance (Supplementary Fig. 1), as well as their correlation with the exonic counts suggest that these molecules represent unspliced precursor mRNAs.

**Figure 1.**
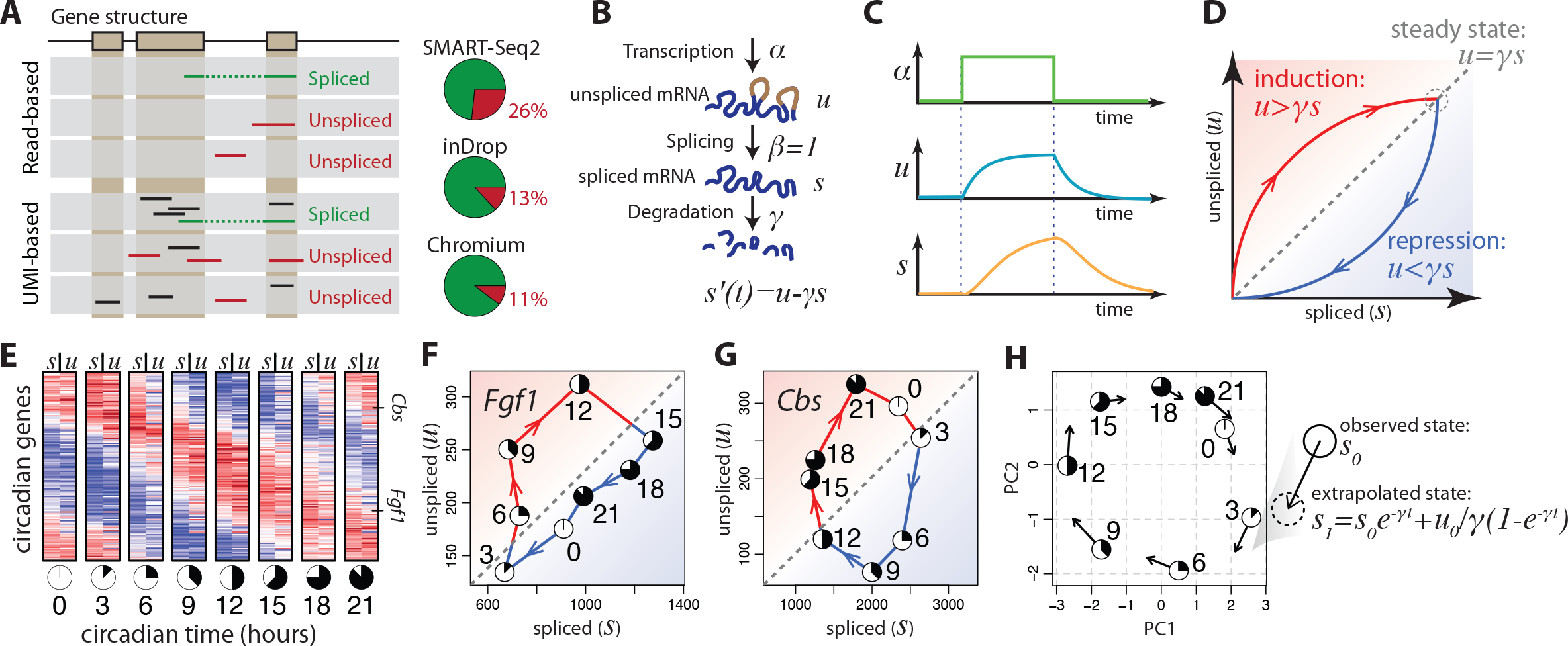
Balance between unspliced and spliced mRNAs is predictive of cellular state progression. **A.** Unspliced mRNA abundance are estimated by separately counting reads that incorporate intronic sequence. In protocols incorporating unique molecular identifiers (UMIs), multiple reads associated with a given molecule can be used to identify unspliced molecules. Pie charts show the fraction of unspliced counts observed for data from different single-cell RNA-seq protocols. **B.** Model of transcriptional dynamics, capturing generation of unspliced pre-mRNAs (*u*) with rate α, its conversion into the mature spliced mRNA (*s*) with rate *β*, and eventual degradation with rate *γ* (scaled in such a way that *β*=1). **C.** Solution of the model in (b) showing pre-mRNA and mature mRNA dynamics in response of changes of the transcription rate α. **D.** Phase portrait showing regions of constant, increasing or decreasing expression with respect to unspliced/spliced mRNA balance. In a steady state condition with a given transcription rate α (dashed circle), the abundance of unspliced mRNA equals the spliced mRNA abundance times the degradation rate γ. Steady states for different transcription rates α fall on the diagonal given by this same γ (dashed line). Levels of unspliced mRNA exceeding that proportion (red) indicate increasing expression of a gene. Similarly, levels of unspliced mRNA below that proportion (blue) indicate ongoing gene repression. **E.** Abundance of spliced (*s*) and unspliced (*u*) mRNAs is shown for circadian-associated genes in a 24h time course of a mouse liver (*15*). At each time point, the expression pattern of unspliced mRNAs was more similar to the spliced mRNA abundance at the next time point than to the current time point, illustrating its power to predict the future expression state. **F,G.** Phase portraits observed for a pair of circadian-driven genes: *Fgf1* (F) and *Cbs* (G). The circadian time of each point is shown using a clock symbol (see bottom of Fig. 1E). The dashed diagonal line shows steady-state relationship, as predicted by a γ fit. *Fgf1* shows continuous increase in expression abundance (*s*) from CT3, reaching its peak expression between CT12 and CT15, followed by continuous decrease in the rest of the cycle. This dynamics is matched by levels of unspliced mRNA (*u*) above and below the steady-state line, respectively. *Cbs* shows similar dynamics, with expression peaks between CT0 and CT3. **H.** Change in expression state at a future time *t*, as predicted by the model, is shown in the space of the first two principal components, recapitulating the progression along the circadian cycle.

To quantify the time-dependent relationship between precursor and mature mRNA abundance, we assumed a simple model for transcriptional dynamics (*15*), where the first derivative of the spliced mRNA abundance (RNA velocity) is determined by the balance between production of spliced mRNA from unspliced mRNA, and the mRNA degradation (Fig. 1B). In a steady state, where a cell is maintaining a constant transcriptional state, RNA velocity is zero, leading to a fixed-slope relationship between the abundance of unspliced (*u*) and spliced (*s*) mRNA molecules: *u* = *γs* (see Methods). During a dynamic process, an increase in the transcription rate *α* results first in an increased abundance of unspliced mRNA, followed by spliced mRNA (Fig. 1C) until a new steady state is reached. Similarly, a drop in the rate of transcription initially leads to a drop in the abundance of unspliced mRNA, followed by spliced mRNAs after a time lag. During induction of gene expression, unspliced mRNAs are present in excess of the expectation based on the degradation rate *γ*, whereas the opposite is true during repression (Fig. 1D). As a consequence, the balance of unspliced and spliced mRNA abundance is an indicator of the future state of mature mRNA abundance, and thus the future state of the cell.

To illustrate that such a simple model can be used to extrapolate the mature mRNA abundance into the future, we examined a timecourse of bulk RNA-seq measurements of the mouse liver circadian cycle (*21*). Unspliced mRNA levels of genes associated with the circadian cycle in this dataset showed a consistent phase advance relative to the levels of mature mRNA (Fig. 1E). Estimating the steady-state unspliced/spliced slope γ based on all time points, we found that many circadian-associated genes showed the expected excess of unspliced mRNA relative to slope *γ* during up-regulation, and a corresponding deficit during down-regulation (Fig. 1F-G).

During dynamic processes, cells move through a high-dimensional expression space determined collectively by all genes. In order to determine if combined extrapolations of all genes could be used to predict future expression state, we solved the proposed differential equation model for each gene and extrapolated the expression state for each measurement throughout the circadian cycle. Plotting both current and extrapolated expression states in the space of the first two principal components, we found that the extrapolated expression state of each sample accurately captured the expected direction of progression of the circadian cycle (Fig. 1H; note how each arrow points in the direction of the next timepoint, and how the tip of the arrow is closer to the next state than to the present one).

Next, to demonstrate our ability to predict transcriptional dynamics in single-cell measurements, we analyzed recently-published single-cell mRNA-seq data on mouse chromaffin cells (*22*) (Fig. 2). During development, a substantial proportion of chromaffin cells, which are neuroendocrine cells of the adrenal medulla, arise from Schwann cell precursors from the neural crest lineage, providing a convenient test case in which the direction of differentiation can be validated by lineage tracing. The transcriptional profiles of these cells were measured using SMART-seq2, which provides high sensitivity and full-length gene coverage (*16*). Phase portraits of many genes induced and repressed during differentiation showed the expected deviations from the predicted steady-state relationship (Fig. 2B-C), though to compensate for the sparseness of the single-cell data we pooled the reads across *k* = 5 neighboring cells. We found that RNA velocity estimates of the individual cells accurately recapitulated the transcriptional dynamics within this dataset, including general movement of the differentiating cells towards chromaffin fate (Fig. 2D), as well as rapid movement of maturing chromaffin cells away from the intermediate state of the differentiation bridge (Supplementary Fig. 2). The RNA velocity estimates also captured cell cycle dynamics involved in the chromaffin differentiation, with the cyclical pattern being apparent in both unbiased PCA projection, as well as focused analysis of cell-cycle associated genes (Supplementary Fig. 2).

**Figure 2.**
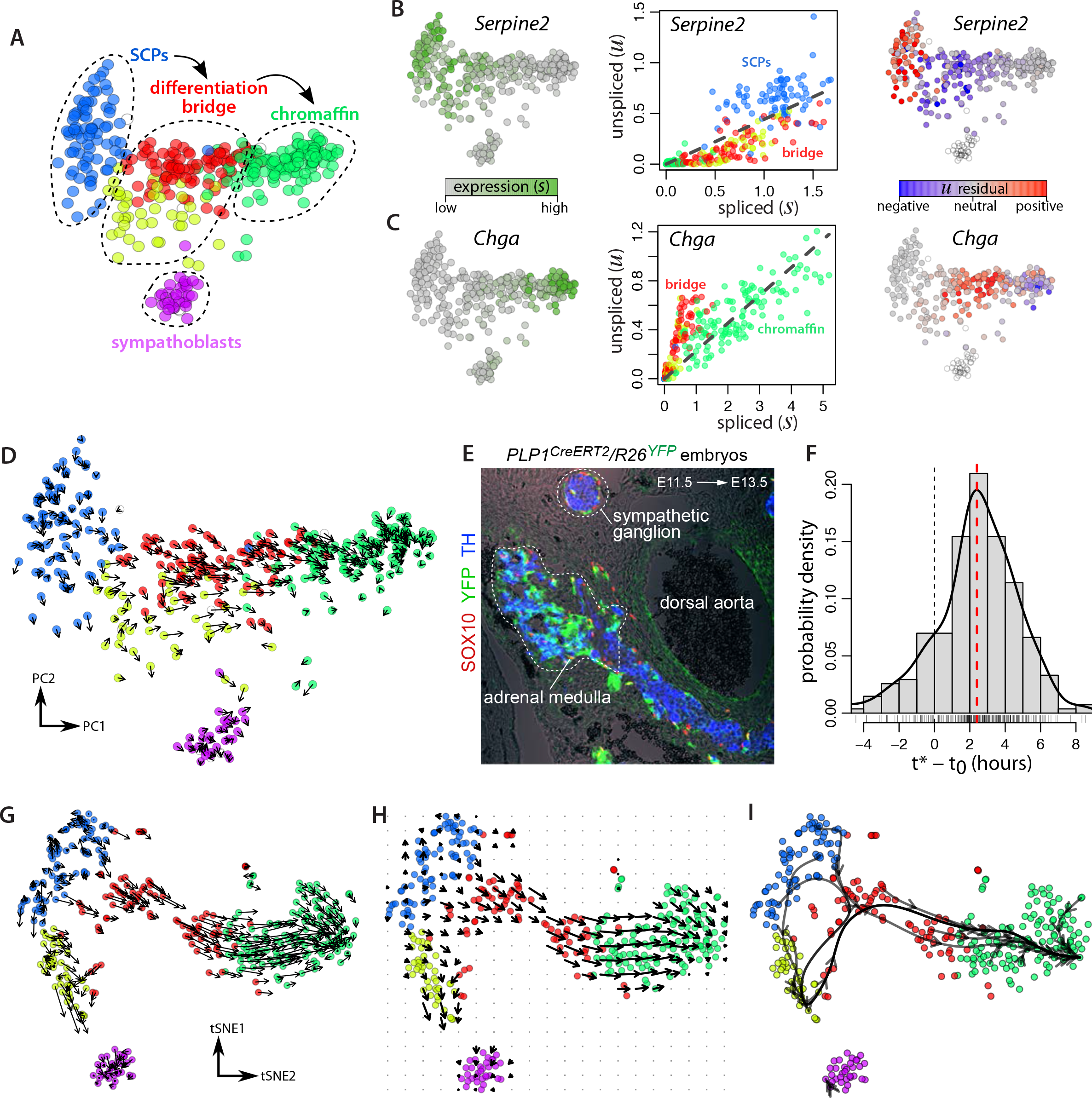
RNA velocity accurately recapitulates dynamics of chromaffin cell differentiation. **A.** A PCA projection shows major subpopulations associated with differentiation of Schwann cell precursors (SCPs) into chromaffin cells in E12.5 mouse. **B,C.** Expression pattern (left), unspliced/spliced phase portraits (center), and *u* residuals (right) are shown for two example genes: *Serpine2* (B) which is repressed and shows bridge unspliced mRNA levels below the steady-state fit, and *Chga* (C) which is induced during differentiation and shows unspliced mRNA levels above steady-state slope in the bridge cells (red, yellow). Each point represents a cell, colored according to its cluster membership as in (a). The read counts were pooled across *k*=5 nearest cell neighbors. **D.** Projection of the observed state and extrapolated future state (arrows) shown on the first two principal components (PCs). RNA velocity was estimated from gene-relative *γ* fits for the individual cells (*i.e.* without cell or gene neighborhood pooling). **E.** SCP-to-chromaffin cell transition as evidenced by the lineage tracing with SCP-specific PLP1-CreERT2 line. A cross-section through the developing adrenal medulla region including dorsal aorta, sympathetic ganglion and a large cluster chromaffin cells in E13.5 mouse embryo are shown. Note high proportion of TH+/YFP+ cells in the developing medulla as compared to the absence of such double-positive cells in the sympathetic ganglion. **F.** Estimation of the velocity extrapolation time lag. A distribution of differences between the observed position of a cell in a chromaffin differentiation timecourse (*t*_*0*_) and the position showing best match to the extrapolation of that cell (*t*\*) is shown (see Supplementary Fig. 7 for details of the estimation). The mode of the distribution was 2.1 hours (red vertical line). **G.** Visualization of the velocities on a pre-defined low-dimensional cell embedding. The velocities are projected on the t-SNE embedding of the E12.5 chromaffin data from the original publication (*22*), using local transition probabilities based on correlation between the velocity and expression state differences of neighboring cells (see Methods). Velocity estimates based on cell kNN count pooling (*k*=5) was used. **H.** Same velocity field is visualized using a Gaussian smoothing on a regular grid. **I.** Predicted cellular trajectories, as modeled by a Markov process with transition probabilities determined based on the velocity estimates (see Methods). Trajectories were simulated for each cell and clustered into 10 clusters. The centroid trajectories of each cluster are shown, using spline smoothing.

Several adjustments can be used to enhance RNA velocity estimation, reducing impact of single-cell measurement noise and gene-specific aberrations. The robustness of the RNA velocity estimates can be improved by pooling of transcript counts across *k* most similar cells (Supplementary Fig. 2). Similarly, pooling of counts can be performed across well-correlated genes, based on the assumption that such genes are also subject to the same up-/down-regulation dynamics (Supplementary Fig. 2). Velocity estimates can also be refined by estimating the contribution of extraneous transcripts to the counts of different genes through analysis of molecules spanning both exonic and intronic regions (Supplementary Fig. 3) or, in case of pooled signal, by fitting *γ* on highest and lowest quantiles of expression (Supplementary Fig. 2).

The gene-relative estimation of *γ* in the analysis presented above relies on the ability to observe a given gene around its steady-state somewhere within the measured dataset. Such estimates cannot detect components of RNA velocity driven by genes that are observed strictly outside of their steady state, such as chromaffin maturation genes up-regulated at the very end of the observed differentiation, or neural-crest genes that are already being actively down-regulated in the initial Schwann cell precursor stage (Supplementary Fig. 4). To address this limitation we developed a structure-based model to predict the steady-state relationship between spliced and unspliced RNA based on the structural parameters of the genes, such as the number of expressed exons and intronic length (Supplementary Fig. 4). The resulting velocity estimates corrected the underestimation at extremes of the chromaffin differentiation trajectory.

As RNA velocity is a vector in high-dimensional expression space, a variety of techniques can be used to visualize the velocity estimates in low dimensions. The observed and extrapolated cell states can be jointly embedded in a common low-dimensional space (e.g. PCA in Figure 2d, t-SNE in Supplementary Fig. 5). Alternatively, velocities can be projected onto existing low-dimensional embeddings based on the similarity of the extrapolated state to other cells in the local neighborhood (Fig. 2G, see Methods). In large datasets, it is easier to visualize the prevalent pattern of cell velocities with smoothed vector fields (Fig. 2H). It is important to note that velocity estimation does not assume existence of an underlying field potential, which the cell flow is expected to minimize (*13*, *14*). In this sense, the RNA velocity is a strictly more general representation than the Waddington landscape, and in contrast to the latter can capture complex situations such as cyclical trajectories or opposing flows (Supplementary Fig. 2).

Cell-specific RNA velocity estimates provide a natural basis for quantitative modeling of cell fates. However, as extrapolation of the expression state is based on its first derivative (velocity), it shows increasing deviation from the true expression manifold at longer time scales. Based on the EdU pulse incorporation data in adrenal medula (Supplementary Fig. 6), we estimate that the velocity-based extrapolation is able to predict cellular state 2.5-3.8 hours into the future (Fig. 2F, Supplementary Fig. 7). A predictive time-scale of several hours is also consistent with the ability to resolve cell-cycle events. Given that the extrapolation is linear, this time-scale depends on the curvature of the expression manifold, with simpler linear manifolds allowing for longer extrapolation times. Prediction of cell fates on curved manifolds at longer time scales can be approached under the assumption of ergodicity. For instance, cell trajectories can be modeled as a Markov process with transition probabilities determined by the local velocity field (Fig. 2I).

To confirm that our approach was able to capture differences in the magnitude of transcription velocity, we examined another mouse chromaffin differentiation dataset, taken at a later developmental time point (E13.5). The resulting velocities recapitulate chromaffin differentiation in a way similar to the earlier time point (Supplementary Fig. 8), however significantly lower magnitude of velocities was reported for the cycling part (yellow) of the chromaffin bridge (Supplementary Fig. 6). We quantified the relative abundance of *Sox10*+ Schwann cell precursors, *Htr3a*-GFP+ bridge cells, and *Th*+ chromaffin cells in tissue sections. Indeed we found that the developmental dynamics of chromaffin cell production slowed down at E13.5 as compared to E12.5 based on the ratio of progenitors and resulting TH+ cells, consistent with lower predicted velocity (Supplementary Fig. 6).

To demonstrate the generality of our approach we analyzed data generated using other single-cell RNA-seq protocols. We found that velocity estimates accurately recapitulated dynamics of macrophage maturation in mouse bone marrow measured using the in Drop protocol (Supplementary Fig. 9) and hippocampus development measured using 10X Genomics Chromium (below). Based on these results, we believe that most single-cell mRNA-seq protocols can be used to estimate RNA velocity.

We next applied RNA velocity to a branching lineage of the developing mouse brain. The mammalian nervous system develops from proliferating radial glia cells that first generate a variety of neuronal cell types, then astrocytes and ependymal cells, and finally oligodendrocyte precursor cells, which in turn proliferate and give rise to the oligodendrocyte lineage. In the hippocampus, neurogenesis occurs late, and therefore it is possible to observe the branching of all major brain lineages simultaneously. We estimated RNA velocity for a total of 18,213 cells (postnatal day 0: 8,113 cells; postnatal day 5: 10,100 cells). In this dataset, 6,088 genes showed detectable levels of intronic reads. After removing vascular and immune cells, and GABAergic and Cajal-Retzius neurons (which originate from outside the hippocampus), t-SNE embedding revealed a complex lineage with multiple branches (Fig. 3A, Supplementary Fig. 10). We identified the tips of the branches as corresponding to astrocytes, oligodendrocyte precursors (OPCs), dentate gyrus granule neurons, and pyramidal neurons of the five fields of the hippocampus: the CA1, CA2, CA3, hilus and subiculum (Supplementary Fig. 10).

**Figure 3.**
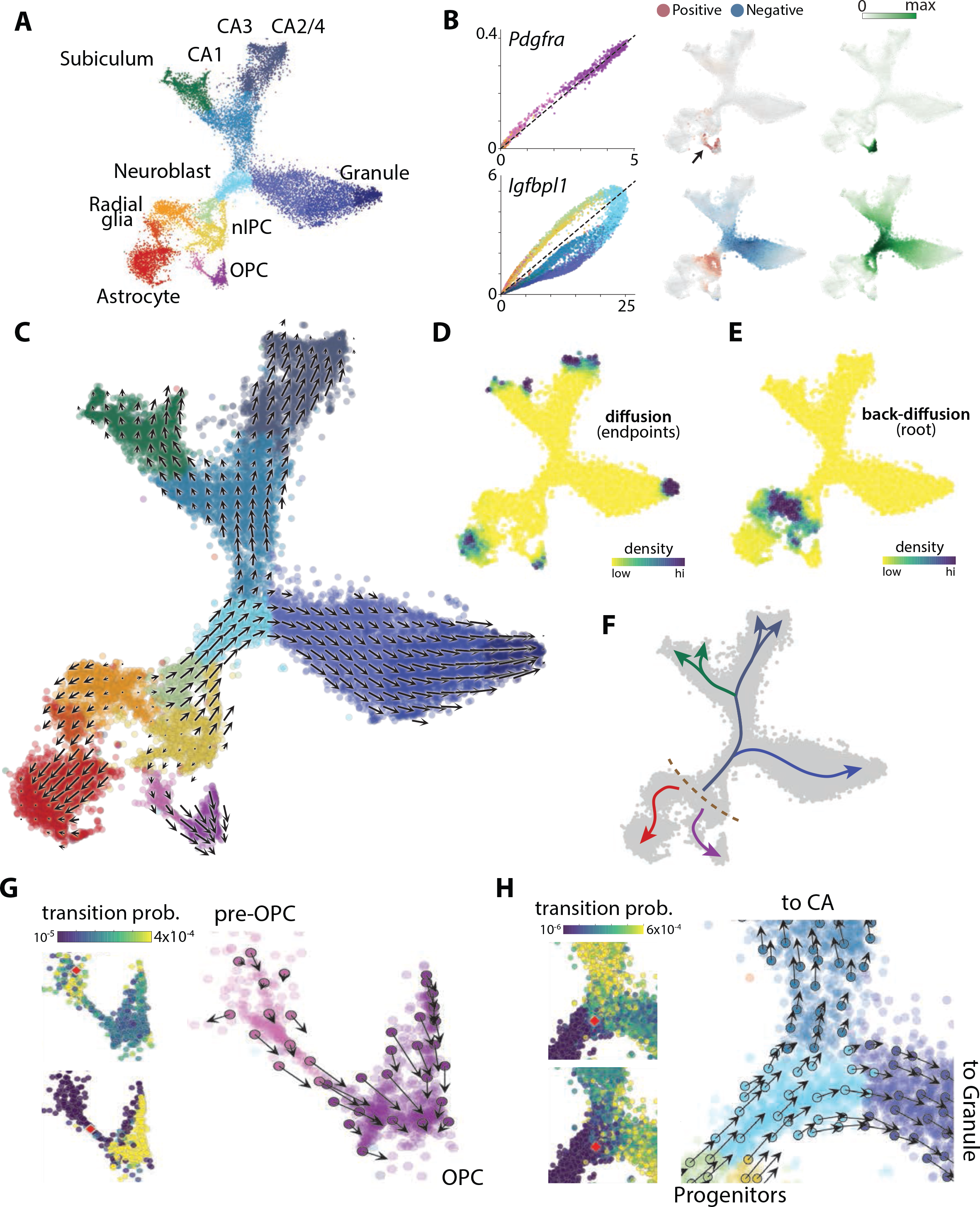
RNA velocity field divergence describes fate decisions of major neural lineages in the hippocampus. **A.** A t-SNE embedding of the developing hippocampus dataset, colored by major subpopulations associated with both transient and mature states. **B.** Dynamics unspliced and spliced RNA over the expression manifold shown for two genes. Left, phase portrait colored as in (A). Middle, RNA velocity field. Right, expression profiles. Velocity and expression was obtained by pooling over nearest neighbours (Methods). **C.** Velocity field associated to the two dimensional t-SNE embedding. Arrows show the local average velocity evaluated on a regular grid. **D.** Differentiation endpoints identified as high density regions on the manifold after a forward Markov process using the velocity-based transition probabilities. **E.** Root of the branching tree identified using the same Markov process in reverse. **F.** Summary schematic of the RNA velocity field shown in (C - E). **G.** Commitment to oligodendorcytic fate. Left, visualization of single-step transition probabilities from two starting cells (red) to neighbouring cells. Right, single-cell velocities of a random subset of cells shown on the t-SNE embedding in (C). **H.** Fate decision of neuroblasts. Left, visualization of single-step transition probabilities from two starting cells (red) to neighbouring cells. Right, single-cell velocities of a random subset of cells shown on the t-SNE embedding in (C).

Phase portraits of individual genes showed specific induction and repression of gene expression along the manifold (Fig. 3B, Supplementary Fig. 12). For example, the expression of *Pdgfra* (a marker of OPCs) was induced in pre-OPCs and then maintained in OPCs; it showed a corresponding positive velocity in the pre-OPC state, but neutral in the OPCs. Similarly, *Igfbpl1* was expressed specifically in neuroblasts, and showed a positive velocity from radial glia to neuroblasts, but a negative velocity going from neuroblasts to the two main neuronal branches.

The RNA velocity vector field showed a strong directional flow towards each of the main branches (Figs. 3C, Supplementary Fig. 11). At the apparent root state, a small group of cells arranged in a band (Fig. 3F, dashed line) lacked velocity, and were surrounded on each side by cells with velocities pointing outwards in opposite directions. We identified these cells as radial glia based on the expression of markers including *Hes1* and *Hopx* (Supplementary Fig. 10). Radial glia are very likely the true origin of the lineage tree of the hippocampus (*23*), thus demonstrating the power of RNA velocity to directly identify the actual root state.

Having a velocity field across the manifold further allowed us to use computational methods to automatically and unambiguously determine both the root and the terminal states. Using a Markov random walk model on the velocity field, the terminal states were identified as sinks (Fig. 3D), and the single root state as the source of velocity field diffusion, demonstrating our ability to orient the lineage tree without prior knowledge about the developmental process. About half of the root cells were in mitotic phase (Supplementary Fig. 10B), suggesting a population of cells undergoing intermittent cell division to maintain a stem cell pool.

On one side, RNA velocity pointed to differentiation into astrocytes (expressing *Aqp4*) without intervening cell division. Radial glia alternatively entered a pre-OPC state, leading through a narrow passage to proliferating OPCs. We speculated that the narrow passage represented the actual moment of commitment to the oligodendrocyte lineage. The Markov process model allowed us to quantify fate choice probabilities at specific points on the manifold. At this microstate level, fate choice is likely a non-deterministic process involving the tilting of gene expression in favor of one or the other fate, followed by a lock-in of the final fate once transcription factor feedback loops are established (*24*). Comparing the probability distribution of future states for a cell starting among the pre-OPCs, versus one starting in the narrow passage leading to OPCs, revealed a clear difference, where the latter cell was overwhelmingly likely to end up as a fully formed OPC, whereas the former was as likely to remain in the pre-OPC state (Fig. 3G).

Some cycling cells expressed neurogenic transcription factors (*e.g*. *Neurod2*, *Neurod4*, *Eomes*) and those cells showed velocity pointing toward an immature neuroblast state, leading towards the three main neuronal branches in the upper part of the manifold. Granule neurons of the dentate gyrus first split from the hippocampus proper, and a second split divided the hippocampal cells into subiculum/CA1 and CA2-4, respectively (Supplementary Fig. 11), in agreement with the major functional and anatomical subdivisions of the hippocampus. Interestingly, the subiculum shared a branch with the CA1, whereas anatomically the CA1 is typically considered more closely related to the CA2, 3 and 4. Note however that the tissue we examined was young, and the ultimate fate of cells along each branch will have to be established in experiments that span the full maturation of the hippocampus to the adult stage.

The detailed, single-cell view of a branching lineage allowed us to ask questions about fate choice. For example, examining two adjacent neuroblasts, just at the entrance to the branching point between CA and granule fates (Fig. 3H), we found that although their current states were nearest neighbors (in gene expression space), their futures were already tilted towards different fates.

To determine if RNA velocity could be detected in the human embryo, we performed droplet-based (10X Chromium) single-cell mRNA-seq of the developing human forebrain (10 weeks post-conception). We computationally identified and isolated the glutamatergic neuronal lineage (Fig. 4A), and calculated RNA velocity as before. Reassuringly, we found a strong velocity pattern originating from a proliferating progenitor state (radial glia), and proceeding through a sequence of intermediate neuroblast stages to a more mature differentiated glutamatergic neuron expressing *Slc17a7* (the vesicular glutamate transporter used in forebrain excitatory neurons). We used principal curve analysis to order the cells according to differentiation pseudotime, and examined temporal progression of transcription in human primary cells. As expected, we found that unspliced RNAs consistently preceded spliced mRNAs during both up-and down-regulation (Fig. 4B). Interestingly, we observed both fast and slow kinetics. For example, *RNASEH2B* showed fast kinetics, with little difference between unspliced and spliced RNAs. In contrast, genes such as *DCX, ELAVL4* and *STMN2* showed evidence of an initial burst of rapid transcription, followed by sustained transcription at a reduced level (as evidenced by the shape of the unspliced RNA curve, Fig. 4C), with spliced transcripts following a noticeably delayed trajectory. Such dynamic induction with overshooting has been proposed to help quickly induce genes whose degradation kinetics are slow (*15*), but have not been possible to study in human embryos. It also suggests that a more detailed modeling of gene-specific transcriptional kinetics should improve the RNA velocity estimates.

**Figure 4.**
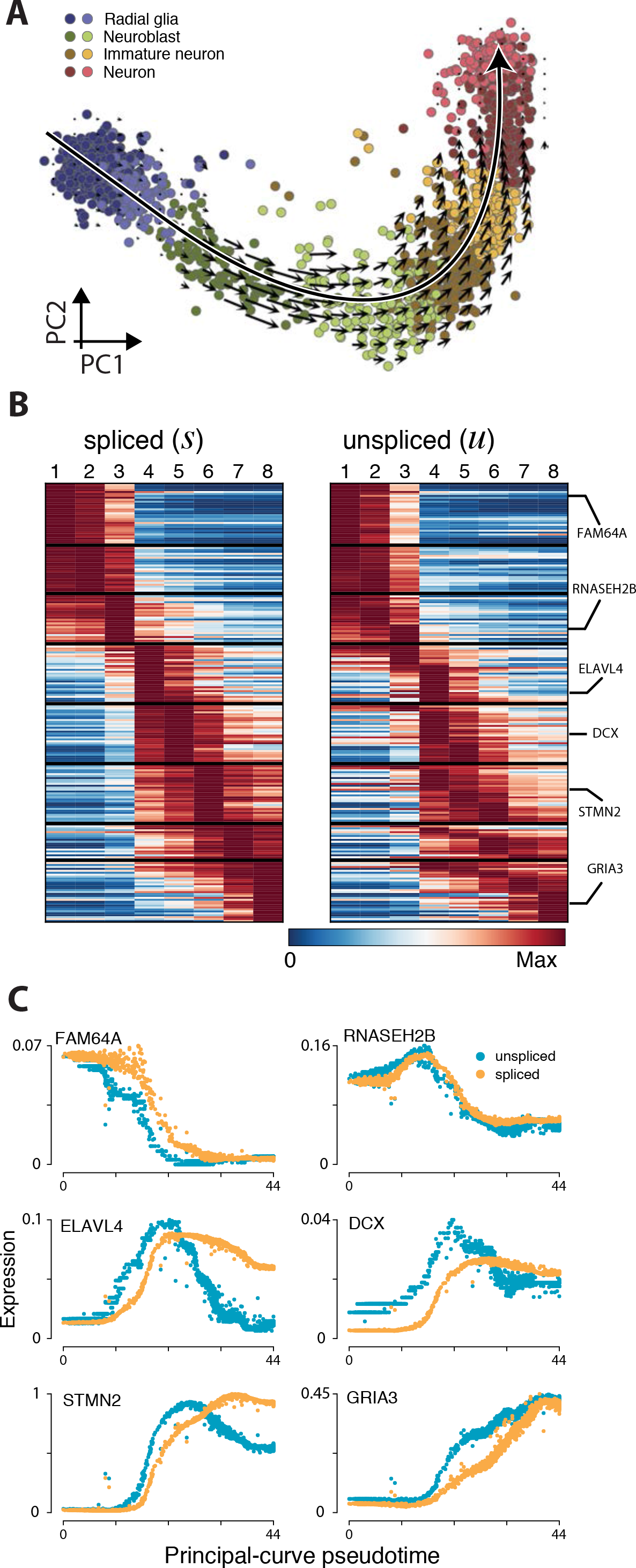
Kinetics of transcription during human embryonic glutamatergic neurogenesis. **A.** PCA projection of human glutamatergic neuron differentiation at post-conception week 10, shown with velocity field. Colors indicate cell types and intermediate states. A principal curve passing through the datapoints, and oriented by the velocity field directions, is indicated by the long arrow. **B.** Unspliced expression anticipates spliced expression. Dynamic genes were sorted by their spliced-count maxima and unspliced-count maxima, and clusters were sorted by directional pseudotime according to the principal curve. **C.** Examples of pseudotime series showing dynamics of pre-mRNA and mRNA expression for six genes involved in glutamatergic neuron maturation. Unspliced traces were rescaled through multiplication by *γ*.

RNA velocity reveals the temporal dynamics of single-cell gene expression on a timescale of hours, matching the unfolding of developmental, regenerative and reactive processes in both human and other mammals. It reveals the landscape of molecular states in great detail, and provides local velocity vectors that can be used to model commitment, fate choice and the precise kinetics of transcription *in vivo*. RNA velocity enables the detailed study of dynamic processes in complex tissues and organs, and will greatly facilitate lineage analysis in the human embryo.

## Supplementary Figure Legends

**Supplementary Figure 1. Correlation of the total spliced and unspliced counts.** The scatter plots (A-C) show the relationship between the total abundance of spliced (*s*) and unspliced (*u*) counts for **A.** Chromaffin E12.5 data, measured by SMART-seq2, **B.** Mouse bone marrow data measured using in Drop, and **C.** Mouse dentate gyrus dataset measured using 10x Chromium platform. log_10_(count +1) values are shown. Pearson linear correlation coefficient is shown in the upper left corner. All genes with non-zero total *s* and *u* counts and intronic length greater than 100bp are shown.

**Supplementary Figure 2. PCA visualization of chromaffin E12.5 velocities. A-D.** Velocity estimates on the chromaffin E12.5 dataset are shown by projecting both observed and extrapolated cells onto the space of first five principal components. The velocity was estimated using gene-relative fit for individual cells (*i.e*. without cell or gene pooling). Overall, PC1 captures the main chromaffin differentiation axis, PC2 captures separation between sympathoblasts and other cell types, PC3 separates bridge-specific (red) cells from others, PC4 and PC5 together capture cell cycle signature of the cycling bridge cells (yellow) – as seen in the last (PC4 *vs*. PC5) panel. **E.** Gene-relative velocity estimates are shown with *k*=5 cell kNN read pooling. **F.** Velocities estimated pooling reads across neighboring cells (*k*_*cells*_=5) and well-correlated genes (*k*_*genes*_=20). **G.** Velocity estimates, with γ slope and offset fit using only cells within the top/bottom 2% expression quantile of each gene. As such approach works robustly on smoothed data, cell kNN pooling (*k*_*cells*_=5) was used in calculating the estimates. **H.** To emphasize cell cycle trends, velocity estimates from the previous panel were subset to include only cell cycle-related. The genes were selected using GO annotations. The resulting observed and extrapolated states were visualized by projecting on the first two PCs. Sympathoblast cells, which also undergo cell cycle within the dataset, were excluded from this visualization.

**Supplementary Figure 3. Fitting offset of intronic read counts using spanning reads.** Fitting of non-specific unspliced count offset using spanning read counts is shown for five example genes (rows). Gene name is given in the lower right corner of each plot. For each gene (row), the first column (**A**) shows expression (spliced count abundance) of the gene using tSNE layout (see Fig. 2D of the main manuscript). The second panel (**B**) shows a scatter plot illustrating the observed dependence between the spanning (x axis) and intron-only (y axis) read counts. The dashed line shows the regression fit that is used to determine the y axis intercept (intronic read count offset). The third panel (**C**) shows relationship between exonic and intronic (intron-only) counts. The dashed black line shows a gamma fit using the intronic count intercept determined from B, and grey dashed line uses zero intercept. The fourth column (**D**) shows unspliced count residuals (basis of the subsequent velocity estimates) calculated using spanning-read based offset from B. The last column (**E**) shows residuals calculated using default zero-offset. Genes with high offset values were chosen as examples

**Supplementary Figure 4. Structure-based velocity estimation.**

**A,B.** For genes that are observed only outside of the steady state, such as genes upregulated late in the chromaffin differentiation (A) or down-regulated early in the SCPs (B), gene-relative γ fit will likely deviate from its steady-state value.

**C,D.** To correct for such effects, a structure-based *γ* fit will first predict *γ* for every gene based on its structural parameters, and then use *k* most correlated genes in the dataset to adjust M-value (*M* = log_2_[*u*_o_/*u*_*ss*_], where *u*_*ss*_ is the unspliced counts predicted from spliced counts under steady-state, and *u*_o_ is the observed unspliced count) using robust mean, and re-estimate *γ*.

**E.** Scatter-plot comparing gene-relative and structure-based *γ* estimates, with colored circles highlighting *γ* adjustments for genes down-regulated early in SCPs (blue) and up-regulated late in chromaffin cells (green). The values are shown on a natural log scale.

**F-I.** RNA velocity in the chromaffin E12.5 dataset, based on the structure-based *γ* estimates, shown on the first five PCs.

**Supplementary Figure 5. Joint t-SNE visualization of observed and extrapolated chromaffin E12.5 cells.**

**A.** The chromaffin E12.5 velocities estimated using gene-relative fit, with *k*=5 cell pooling are shown by joint embedding of observed (circles) and extrapolated cells (end of arrows) using t-SNE.

**B.** Analogous joint t-SNE embedding for the chromaffin E12.5 velocities estimated using structure-based model.

**Supplementary Figure 6. Analysis of developmental dynamics in early adrenal medulla.**

**A.** Changes in numbers of Sox10+ SCPs (progenitors), Htr3a-GFP+ bridge cells (intermediate progenitors) and TH+ chromaffin cells during subsequent developmental stages. Note that at E13.5 the pool of bridge cells decreases as compared with E12.5, while Sox10+ progenitors keep increasing their numbers.

**B.** Proportion of Sox10+ progenitors over generated TH+ chromaffin cells per next developmental stage (TH+ cells accumulated at the previous developmental stages were subtracted). Note that bigger numbers of Sox10+ progenitors generate proportionally less TH+ chromaffin cells at E13.5 as compared to E12.5.

**C.** The barplots compare the ratio of total unspliced and spliced mRNA molecules between E12.5 (solid bars) and E13.5 (shaded bars) time points. Statistically significant (p<10^−5^) decrease in the unspliced/spliced molecule count ratio is observed for the cycling subpopulation (yellow) of the chromaffin bridge at E13.5, indicating lower RNA velocity.

**D.** Immunohistochemistry on transversal sections of mouse embryos at the level of developing adrenal medulla. Dotted line outlines developing medulla and cortex at E11.5.

**E.** Measurements of EdU incorporation and retaining in various populations of adrenal medulla 14 and 24 hours after the single pulse. The analysis stage is E13.5. Note that first TH+ cells that retain both weak GFP and EdU (immediate progeny of Htr3a-GFP bridge cells) are identified in the tissue 14 hours after EdU injection. Their numbers are comparable to the proportion of EdU-labeled Sox10+ progenitors 24 hours after EdU pulse. Yellow arrowheads in immunohistochemistry panel point at GFP-retaining TH+ cells.

**F.** Schematic explanation of differentiation progression in chromaffin cell lineage. Note that Sox10+ SCPs proliferate strongly. At the same time, very few independently dividing cells were detected in more mature GFP−/TH+ population of chromaffin cells 4 hours after EdU pulse (data not shown).

Supplementary Figure 8. RNA velocity is can predict cellular trajectories on the timescale of hours.

**A.** PCA projection of E12.5 dataset showing, as a reference, major subpopulations in the chromaffin differentiation (same as in Fig. 2A of the main manuscript).

**B.** Optimal extrapolation distance along the chromaffin differentiation trajectory. The plots show correlation between the velocity vector and cell expression difference vector (y axis) for the cells ordered by chromaffin differentiation pseudotime (x axis). Correlation profiles for three example cells are shown, with pseudotime of each cell (*t*_*0*_) and pseudotime of the maximal correlation (*t*\*) marked by the black and red dashed lines, respectively.

**C.** The optimal extrapolation distances (from *t*_0_ to *t*\*, x axis) are shown for all of the cells along the chromaffin differentiation pseudotime (y axis). The distribution of these distances is shown in the Fig. 2F of the main manuscript. The cells at the extreme of the pseudotime (beyond the 10% thresholds marked by vertical dashed lines on the current plot) were excluded, as estimation of pseudotime within such extremes is not expected to be robust. For the Fig. 2F of the main manuscript, the pseudotime time differences were translated into real hours, based on the 14 hour total chromaffin differentiation time (Supplementary Fig. 6).

**Supplementary Figure 8. PCA visualization of chromaffin E13.5 velocities using estimated using gene-relative model.** PCA projections are used to show E13.5 chromaffin dataset velocities, as estimated by the gene-relative model with *k*=5 cell kNN pooling. Projections onto the first five PCs are shown. The cell clusters are colored using the same color scheme as for E12.5 dataset (see Fig. 2A of the main manuscript).

**Supplementary Figure 9. Velocity estimates in mouse bone marrow cells measured using in Drop protocol.**

**A.** Grid visualization shows RNA velocity estimates for the in Drop mouse bone marrow dataset on a t-SNE embedding.

**B.** Major cell populations are labeled based on manual annotation. The velocity flow in (A) captures macrophage maturation, starting from the dividing cells on the right, all the way to macrophages showing Il1b activation on the right.

**C.** Expression profiles for five marker genes are shown on the t-SNE embedding.

**D-G.** The plots illustrate gene-relative model fits for several example genes. The first column (D) shows spliced molecular counts in different cells. The second column (E) shows unspliced molecular counts. Third column (F) shows phase portrait of a gene (unspliced vs. spliced dependency) and the resulting *γ* fit (dashed red line), as determined using extreme quantile method. Each point corresponds to a cell, colored according to cluster labels shown in (B). The last column (G) shows unspliced count signal residual based on the estimated *γ* fit, with positive residuals indicating expected upregulation, and negative residuals indicating expected downregulation of a gene.

**Supplementary Figure 10. Branching developmental trajectories of developing hippocampus.**

**A.** t-SNE embedding of the developmental dentate gyrus dataset. Cells are colored by cluster identities, with labels shown for the major cell types.

**B.** Expression of radial glia (and astrocyte) marker *Hes1,* and cell cycle genes *Top2a* and *Cdk1* shown on the t-SNE embedding.

**C.** Marker genes of different regions of the hippocampus show prominent expression signals at different extremities of the branching embedding. Scale bars, 0.5 mm.

**Supplementary Figure 11. Single cell velocity estimates for individual cells in the embryonic hippocampus dataset.** Arrows indicate the extrapolated state projected onto the t-SNE embedding of the manifold.

**Supplementary Figure 12. Selected phase portraits and fits of the degradation rate (*γ*) for the developing cells in the embryonic hippocampus dataset.**

For each gene, the first column shows spliced-unspliced phase portrait. Dashed line represents the *γ* fit. The second column shows the magnitude of the residuals (*i.e*. velocity) and its sign for several genes involved in the nervous system development. The third column shows the expression profile of the gene.

## Methods

### Theoretical description of RNA velocity

Based on the model of transcription shown in Fig. 1, we can write down the rate equations for a single gene, which describes how the expected number of unspliced mRNA molecules *u*, and spliced molecules *s*, evolve over time: 
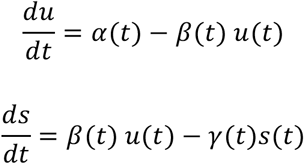

Here, *α*(*t*) is the time-dependent rate of transcription, *β*(*t*) is the rate of splicing, *γ*(*t*) is the rate of degradation. Under an assumption of constant (time-independent) rates *α*(*t*) = *α*, *γ*(*t*) = *γ*, and setting *β*(*t*) = 1 (i.e. measuring all rates in units of the splicing rate), the rate equations simplify to: 
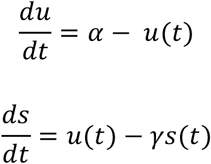

The complete solution to the rate equations is given by:

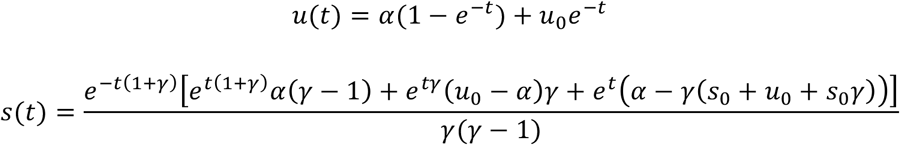
with the initial conditions *u*(0) = *u*_0_ and *s*(0) = *s*_0_. This solution can be used to extrapolate mRNA abundance *s* to a future timepoint *t*_1_, under the assumption stated above, by entering the current state of the cell as *u*_0_ and *s*_0_, and then computing *s*(*t*_1_).

The equations above hold for a single gene. Across all genes, the same equations hold under the same assumptions, but with gene-specific rate constants. Note that setting *β*(*t*) = 1 for all genes implies that we assume a common, constant rate of splicing. The normalized degradation rate *γ* varies among genes and needs to be estimated in a gene-specific manner. In steady-state populations, where *ds*/*dt* = 0, we can determine *γ* of a given gene as the ratio of unspliced to spliced mRNA molecules:

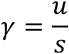

The steady-state assumption may be realistic for genes expressed in populations known to be terminally differentiated. However, for genes expressed transiently during development, or in cases where the terminal population was not sampled, the steady-state assumption will fail. The subsequent sections will detail how *γ* can be estimated without the steady-state assumption.

More problematically, we do not know *α*, nor can it be easily estimated. This prevents us from extrapolating *s*(*t*) into the future. Instead of assuming a constant *α*, we therefore estimate *s*(*t*) using one of two alternative assumptions:

#### Model I. Constant velocity assumption

We assume that for the purposes of *s*(*t*) extrapolation, the rate of change of the spliced molecules remains constant. That is, we assume *ds*/*dt* = *v* is constant, so that the current rate of increase or decrease in spliced mRNA molecules continues into the future. Under this assumption, extrapolation is trivial, since 
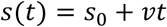

In other words, extrapolation amounts to taking the current number of mRNA molecules and adding the current rate of change multiplied by the extrapolation time step. This assumption works well in practice as long as the time step is short. For longer extrapolation, *s*(*t*) can become negative if *v* < 0 (i.e. in the case of a down-regulated gene). This requires clipping the values at zero.

#### Model II. Constant unspliced molecules assumption

Alternatively, we can extrapolate *s*(*t*) assuming that the number of unspliced molecules stays constant, i.e. that *u*(*t*) = u_0_. This reduces the problem to a single rate equation:
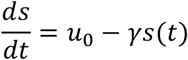

The solution then becomes

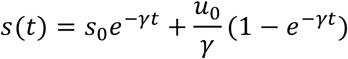

In practice, we found that at short extrapolation timescales both approaches yielded very similar results. We will indicate below when we used Model I or the Model II.

Assuming gene independence, the overall RNA velocity of the cell is a multidimensional vector comprised of the individual gene velocities.

### Estimation framework

In this section we give a description of the analysis framework we used the estimation of RNA velocity and the related data analysis. This analysis logic is implemented separately in R and python environments by velocyto.R and velocyto.py packages, respectively. Parameters, thresholds and other information related to the implementation of each package are described in detail in the next section, and code to reproduce our analysis is available in the companion notebooks at http://velocyto.org.

**Estimation of RNA velocity.** For each gene, the normalized degradation rate *γ* was determined using a least squares fit of the following linear model: *u* ~ *γ* * *s* + o, where *u* and *s* are the size-normalized unspliced and spliced abundances, respectively, observed for given gene across the cells. Specifically, in a given cell *u* = *U/N*, *s* = *S/N*, where *U* and *S* are the number of unspliced and spliced counts, respectively, and *N* is the total number of molecules observed in a given cell. *o* is an offset constant that models baseline intronic counts that might be driven by unannotated transcripts, estimated using one of several possible approaches as detailed further below.

Under Model I, the velocity component v for a given gene in a given cell was assumed to be constant and estimated as *v* = *u* − *γs* − *o*. The extrapolated counts of a given gene in a given cell was then determined as *s_t_* = *max*(0, *s*_0_ + *vt*). Where t is the extrapolation time step, that was chosen such that the total RNA count for each cell did not change substantially.

Under Model II, the displacement of spliced mRNA *Δs* for a given gene in a given cell was estimated assuming constant u as 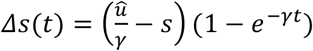, where *û* is the offset-adjusted unspliced count, calculated as *û* = *max*(0,*U* − *o* * *N*)/*N*, and using the default extrapolation time *t* = 1. The extrapolated counts of a given gene in a given cell was then determined as *S*_t_ = *max*(0,S + *Δs*(*t*) *N*), where *S* is the non-normalized spliced count for a gene in a given cell. The normalized extrapolated counts were then calculated as 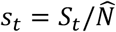, where 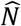 is the extrapolated total size of the cell 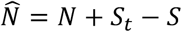.

**Cell nearest neighbor (kNN) pooling.** To improve *γ* estimation, pooling of count data across local neighborhoods was employed. Specifically, cell pooling was implemented by substituting *S*, *U* counts with the sum of the counts from the original cell and its *k* closest cells. k was adapted to match the sparsity and size of the dataset. The estimation of velocity was carried out using pooled count data. The extrapolated state was calculated using the initial (non-pooled) count values. Pearson linear correlation distance on all genes (log scale) was used to find *k* closest cells for the SMART-seq2 datasets, while Euclidean distance in PCA space was used for the larger 10x Chromium and in Drop datasets.

Visualization of cell velocities. The extrapolated state for a cell corresponds to a vector in the same space of the original cell measurement, as such, it can be directly visualized using linear dimensionality reduction approaches such as PCA. For PCA-based visualization (Figure 1H, Figure 2D), principal components were determined based on the observed expression space. The projection of the extrapolated state on the same eigenvectors were then used to position the end of the velocity arrows. For joint t-SNE projection (Supplementary Fig. 5), t-SNE embedding was determined using a union of both extrapolated and observed cells states, using high perplexity (200 in case of Supplementary Fig. 5).

To visualize velocity on an existing embedding (*e.g*. Figure 2G), a transition probability between cells was first estimated. Specifically, transition probability matrix **P** was calculated applying an exponential kernel on the Pearson correlation coefficient between the velocity vector and cell state difference vectors:

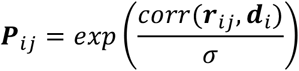
where ***r***_*ij*_ is the difference vector between the expression vectors ***S***_*i*_ and ***S***_*j*_ of cells *i* and *j* transformed with a variance-regularizing transformation 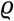 and ***d***_i_ is the 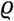-transformed velocity extrapolation vector of the cell *i*:

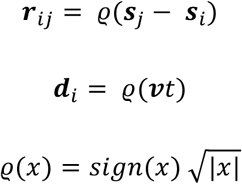

The transition matrix was row-normalized sum so that ∑_*j*_***P***_*ij*_ = 1. Then the transition probabilities *P*_*ij*_ were used as weights to compute a linear combination of the unitary displacement vectors. Given an embedding with the n positions of the cells described by a set of vectors 
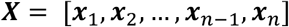
the predicted velocity displacement of a cell was calculated as: 
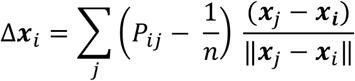
where subtracting *1/n* corrects the estimate for the non-uniform density of points in the embedding.

Visualization of individual cell velocity arrows is not practical for large datasets. For such cases we visualized a vector field showing local group velocity evaluated on a regular grid. The grid vector field was estimated by applying Gaussian kernel smoothing to the velocity vectors of cells around each grid point: 
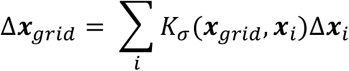
where kernel function *K*_σ_ was defined as: 
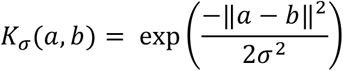

The size of the grid (number of grid points) was chosen depending on the visual scale of the figure.

### Modeling common cell trajectories (Figure 2I, Supplementary Fig. 2.3)

Modeling of cell trajectories was performed based on a Euclidean transition probability matrix, as it provides better control over distant transitions by allowing to explicitly describe the drop off in the transition probability with the increasing expression distance. 
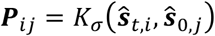
where *s*_0,*j*_ is the (size-normalized) observed spliced expression state of a cell *j*, and ***s***_*t,i*_ is the extrapolated state of the cell *i* at a time *t*. **ŝ** designates a projection of vector ***s*** onto the first 30 principal components. σ = 2.5 was used. Background transition probability capturing the observed cell similarities was calculated as: 
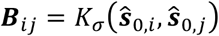

Transition probabilities between cells were restricted to *k*=40 nearest neighbors by setting other values to 0. To correct for local cell density, the rows of each matrix was multiplied by the *diag*(***B***^*b*^). The matrices were row-normalized to unity. The probability of a cell *i* at a discrete time t was estimated as ***P***^*t*^. Most likely position of each cell after *t*_*f*_ = 500 were estimated as the maximum likelihood positions. A trajectory *p*_*i*_ for cell *i* was determined as a path maximizing the total log likelihood: 
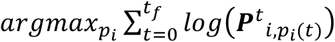
where *p*_*i*_(*t*) is the predicted trajectory position of the cell *i* at a time *t*. To determine prevalent trajectories within a population, individual trajectories *p*_*i*_ were clustered using manhattan distance measuring the difference in the set of cells covered by each path using k-means clustering. 10 clusters were used. The cluster medoids were visualized using xspline smoothing.

### Diffusion start and end-point modeling (Fig. 3)

To find the set of cells that correspond to the differentiation starting point (e.g. early neural progenitors) and end points (*e.g*. neurons and glial cells) we used a Markov process where respectively the transition probability matrix ***T*** or its row sum normalized transpose, where each entry was defined as: 
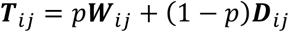
where **W** (after row sum normalization) corresponds to the local Brownian motion component: 
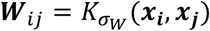
and D (after row sum normalization) represents the local velocity-driven drift: 
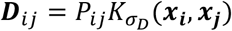

The Gaussian kernel 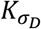 is used to make the diffusion process proceed gradually and locally on the embedding. p<0.5 is the mixture ratio that we set equal to 0.2. Furthermore to avoid the influence of the local density of points on the result, we downsampled the dataset selecting the nearest neighbours of a uniform grid. *σ*_*D*_ was set to the average distance between neighbouring points and *σ*_*w*_ = *σ*_*D*_/2. In both forward and back diffusion we started from a uniform distribution and performed 2500 iterations.

**Pseudotime Analysis.** A principal curve was fit to the subspace consisting of the top four principal components (using the R package princurve). The cell positions were projected onto the curve and the length of the arc between projections was used as pseudotime coordinates. The direction of the pseudotime was determined using the velocity field.

**Gene kNN pooling (Supplementary Fig. 2).** Gene pooling was implemented by pooling counts across k most correlated genes. Gene correlation was assessed using Pearson linear correlation distance on *log*(*s* + 1) values. The resulting matrix represents counts for metagenes. The log ratio of observed to expected unspliced counts was calculated for a given gene and a given cell as 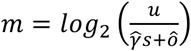, where 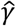 and *ô* are the the slope and offset estimates obtained using metagene matrices. Then, gene *γ* was estimated by taking mean of 2^*log*_2_(*û*/*s*)−*m*^ across all cells with *s* > 0. Under the assumption of constant *u* (Model II), *Δs*(*t*) was then estimated as *Δs*(*t*) = (*s* + *ϵ*)[*e*^−*γt*^(1 − 2^*m*^) + 2^*m*^] − 5, where *ϵ* = 10^−4^.

**Structure-based estimation of RNA velocity (Supplementary Fig. 4)**. Structure-based model starts with the initial estimates (*γ*_*r*_) obtained using gene-relative model described above. Subsequent analysis considered only genes with more than one annotated exon (*n*_*e*_ > 2), with total exonic length *l*_*e*_ over 500bp, and total intronic length *l*_*i*_ over 3kbp. Then a global generalized additive model was constructed to fit the dependency *γ*_*r*_ on the structural parameters of the gene and its expression magnitude: 
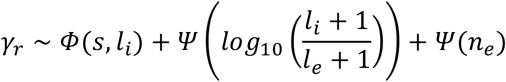
where *Ψ* and *Φ* denote 1D and 2D local smoothing kernels, respectively as implemented by the mgcv package. The model was weighting the observations of each gene by the square root of its total spliced counts across the dataset. The global model was fit omitting genes with disproportionate amount of unspliced counts (likely due to unannotated non-coding transcripts). To do so, total unspliced counts were modeled as ∑_*Cells*_ *u* ~ ∑_*Cells*_ *s* + *l*_*i*_/*l*_*e*_ using a generalized linear model with normal distribution and log link. and genes with pearson deviance exceeding 3 were omitted when fitting the model.

The global model was then used to predict steady-state *γ* (*γ*_*r*_) for all genes passing structural parameter thresholds mentioned above. *γ*_*r*_ was then used to estimate the log ratio of observed to expected unspliced counts as 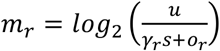, where *o*_*r*_ is the gene specific offset determined as described in the gene-relative model. As we expect *m*_*r*_ estimates of individual genes to be noisy, the next step used k nearest genes to stabilize *m* estimates for each gene.

Specifically for a given gene g, *m* was estimated as a trimmed mean of the *m*_*r*_ values for *k* genes most correlated with gene g across the entire dataset. Default *k*=15 was used, with top and bottom 5 genes trimmed. These robust *m* estimates were then used to estimate RNA velocity as described in “Gene kNN pooling”.

### Analysis pipelines, parameters and implementations details

We implemented the procedures above as two complete pipelines, one in R and one in Python, called velocyto.R and velocyto.py, respectively. These were used to generate all the analyses in the paper, with detailed settings as described in the following sections.

### Details of velocyto.R implementation

**Read annotation.** For Smart-seq2 an implementation counted three types of reads: i) those aligning to an annotated exon (exonic); ii) those aligning to an annotated intron (intronic); and iii) those spanning both an exon and an intron (spanning). As spanning reads are very rare, the subsequent calculations used exonic read count as S, intronic read counts as U. Spanning reads were only used as an alternative method of fitting gene-specific offset *o* (Supplementary Fig. 3). in Drop reads were annotated using dropEst pipeline (*25*), separating molecules into the same three classes. Here, UMIs were used to ensure that a read is annotated is exonic only if all of the reads for that molecule are exonic. Similarly, the molecules were considered intronic only if all of the associated reads were aligned to introns. Molecules with mixed reads were annotated as spanning. However the spanning class was extremely rare in the in Drop data and was not used in the calculations.

**Gene-relative estimation of RNA velocity.** In the basic velocity estimation scheme, *o* was taken to be mean *u* across cells where *s*=0. For the least squares fit, cell-specific regression weights *w* were taken to be *e*^4^ + *s*^4^. Note that *o* and *γ* fit were estimated using a different logic when using quantile fit, spanning-read based fit, or gene *k*NN pooling. To improve stability at low counts, *û* was additionally calculated adding or subtracting one pseudo-count, and a minimal magnitude velocity *v* was reported.

**Cell *k* nearest neighbor (k-NN) pooling.** For the chromaffin data (Figure 2 and related supplements), *k* nearest cells were determined using Pearson linear correlation distance calculated on *log*(*s* + 1) values.

**Gene filtering.** The *γ* coefficients and velocity estimates were calculated for genes meeting a number of filtering criteria: *γ* ≥ 0.1; Spearman rank correlation between *s* and *u* ≥ 0.1; average s counts for a gene ≥ 5 for at least one cell subpopulation (cluster); average *u* counts for a gene ≥ 1 for at least one cell subpopulation; for the datasets where spanning reads were annotated (E12.5, E13.5), average spanning read counts were required to be ≥ 0.5 in at least one subpopulation.

**Extreme quantile fit.** Data smoothed using cell kNN or gene kNN count pooling provided sufficient stability to fit the *γ* coefficient and offset *o* based on the cells at the extreme quantiles of the expression values. Specifically, the linear model *u* ~ *γ* * *s* + *o* for both *γ* and *o* using cells with values of s within the top and bottom 5% for that gene. No regression weights were used.

**Fitting offset using spanning reads.** For Smart-seq2 datasets, the abundance of reads spanning intron and exon boundaries is sufficiently high to enable estimation of the unspliced offset *o*. The offset was estimated using a linear regression

### Details of velocyto.py implementation

#### Annotation of spliced and unspliced reads

For the 10X genomics platform sequencing data was processed using default parameters of the *Cell Ranger* software. We then used the genome annotation (GRCm38.84 and GRCh37.82 from the Cell Ranger pre-built packages) to count molecules while separating them into three categories: “spliced”, “unspliced” or “ambiguous”, as follows. First, we annotated each nucleotide position, keeping track of every exon and intron that overlapped it, taking into consideration e.g. that an intron of one gene model and an exon of another could overlap the same position. Based on this annotation, we then defined exonic and intronic subsections based on the presence of overlapping exons and introns of other genes.

We reasoned that, even in the situation where a subsection was unambiguously annotated as intronic, unspliced counts estimation might be skewed by reads generated as results of antisense transcription of other transcripts or unannotated exons of other transcripts. To reduce the influence of this possible source of noise, already at the level of the counting pipeline, we only considered intron intervals for which there was evidence of the presence and detectability of the unspliced molecule. Specifically, we required the detection of at least one valid read spanning an exon-intron junction in the sequencing library (not necessarily in the same single cell).

For UMI-based technologies, the call for spliced or unspliced was performed molecule-wise. To call for spliced or unspliced molecules the set of reads corresponding to each UMI (cell barcode + random barcode) were taken in consideration as following: if one or more of the reads overlapped with a valid intron then the molecule was considered unspliced; if the previous condition was not satisfied then we marked as exonic a molecule that had at least one read mapping to a exon-exon junction or an unambiguously exonic subsection. The rest of the molecules were annotated as ambiguous and not used in the downstream analyses.

**Extreme quantile fit.** To avoid bias associated with different density of cells in different section of the s vs. u phase diagram and exploiting the large number of cells. We fit gamma parameter by least squares fit considering only the extreme quantiles (top and bottom 2 percentiles) of the sum of spliced and unspliced: s/max(s) + u/max(u). Furthermore, for the genes for which *median*(*u*) > *median*(*s*) (where we usually expect *median*(*s*) ≈ 0.1 * *median*(*u*)), we fixed an upper boundary to the value that gamma could assume to 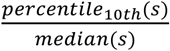. This avoids overfitting the phase portrait when there is evidence that we might not be observing the steady state for a certain gene.

### Dentate Gyrus dataset analysis using velocyto.py (Fig. 3)

The embedding was computed on the correlation similarity matrix using pagoda2 (https://github.com/hms-dbmi/pagoda2). Briefly, gene variance normalization was performed by fitting a generalized additive model of variance on expression magnitude, and rescaling the gene variance by matching the tail probabilities of log residuals from the F distribution to the chi squared distribution with the degrees of freedom corresponding to the total number of cells. Cell distances were determined as 1 − *r*_*ij*_, where *r*_*ij*_ is Pearson linear correlation of the cell *i* and *j* scores on the first 100 principal components of the top 3000 variable genes in the dataset. Clustering was performed using the Louvain community detection algorithm on the nearest neighbor cell graph (*k*=30, pagoda2 implementation). For the velocity analysis lowly expressed (spliced) genes were excluded (requiring 40 minimum expressed counts and detected over 30 cells) and the top 3000 high variable genes were selected on the basis of a non-parametric fit of CV vs. mean (using support vector regression). We kept for the analysis only the 1706 genes that had unspliced molecules above a detection threshold (25 minimum expressed counts and detected over 20 cells). We then normalized the cell total molecule counts to the median. To reduce dimensionality, PCA was performed and the top 20 variable components were selected on the basis of explained variance ratio profile. In this PCA space we built a k-nearest neighbor graph (k=500) based on Euclidean distance and using a greedy balanced k-NN algorithm that limits each node to have no more than 4*k incoming edges. After k-NN pooling we renormalized the molecule counts by the median. Extrapolation was performed using Model I assumptions.

### Human glutamatergic neurogenesis using velocyto.py (Fig. 4)

Pseudotime analysis was performed fitting principal curve on the top four principal components. Clustering was performed determined using Louvain community detection algorithm on the nearest neighbor graph in the same subspace (pagoda2 implementation). For the velocity analysis lowly expressed (spliced) genes were excluded (requiring 30 minimum expressed counts and detected over 20 cells) and the top 2000 most variable genes were selected on the basis of a non-parametric fit of CV vs. mean (using support vector regression). We kept for further analysis the 987 genes that had unspliced molecules above a detection threshold (requiring 25 minimum expressed counts and detected over 20 cells; average spliced counts for a gene 0.06 in a subpopulation and average unspliced counts for a gene 0.007 in a subpopulation). We then normalized the cell total molecule counts to the median. For cell k-NN pooling we built a k-nearest neighbor graph (k=550) based on Euclidean distance in the top six principal components and proceeded as above. After convolution we renormalized the molecule counts by the median and fit the gamma parameter as described above. Genes whose expression peaks at different stages of neurogenesis were selected using a heuristic gene enrichment score: 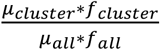 where *μ* indicates the average molecule count of a gene and *f* is the fraction of cells in which the gene is detected. Figure 4B shows the top-enriched genes that showed Pearson linear correlation greater than 0.55 between the estimated velocity and the expression change. Expression change was calculated by dividing pseudotime into 10 equally-sized bins and evaluating finite differences. In Figure 4C we bring spliced and unspliced molecules to a comparable scale, by multiplying spliced molecular counts by the estimated *γ*.

### Human tissue and single-cell RNA sequencing

Human first trimester subcortical forebrain tissue was obtained from elective routine abortions (10 weeks postconception) with the written informed consent of the pregnant woman and in accordance with the ethical permit given by the Regional Ethics Vetting Board (Stockholm, Sweden). Human fetal forebrain tissue was collected and stored in hibernation media with addition of GlutaMAX and B-27 supplements according to the manufacture’s instructions (overnight, 4oC, Hibernate-A, Thermo-Fisher). The tissue was then cut into small cubic pieces of approximately 1-2mm length. Tissue was dissociated using a dissociation kit (Miltenyi, Neural Tissue Dissociation Kit (P)) according to manufacture’s instructions. In short, tissue was prepared in the kit buffer containing 0.067mM beta-mercaptoethanol. After addition of enzyme mix 1 and 2, the tissue was mechanically dissociated using three increasingly smaller gauges of fire polished Pasteur pipettes, pipetted 20, 15 and 10 times up and down respectively. Ultimately, collected cells were stored on ice in PBS containing 1% BSA and immediately prepared for single cell library preparation. Single-cell RNA sequencing was performed using the 10X Genomics Chromium V2 kit, following the manufacturer’s protocol, and sequenced on an Illumina Hiseq 2500.

### Validation of chromaffin differentiation velocity in mouse

#### Mouse lines

For all experiments, the day the plug was detected was considered as E0.5. All animal work was permitted by the Ethical Committee on Animal Experiments (Stockholm North committee) and conducted according to The Swedish Animal Agency’s Provisions and Guidelines for Animal Experimentation recommendations. *Htr3a*^*EGFP*^ animals were received from MMRRC and provided by J. Hjerling-Leffler laboratory (Karolinska Institutet, Sweden) (https://www.mmrrc.org/catalog/sds.php?mmrrc_id=273.

#### EdU incorporation and analysis

14 hrs or 24 hrs prior to embryo collection, pregnant females received an intraperitoneal injection of EdU (50 μg/g of body weight). EdU was visualized using the Click-iT EdU Alexa Fluor 647 Imaging Kit (Life Technologies) according to manufacturer’s instructions.

#### Immunohistochemistry

Immunohistochemistry was performed as previously described (*22*). Briefly, embryos were collected and fixed in 4% paraformaldehyde in PBS (pH 7.4) at 4°C for 5 hours. Samples were washed in PBS at 4°C for one hour and cryoprotected by incubating at 4°C overnight in 30% sucrose in PBS. Tissue samples were subsequently embedded in OCT and frozen at – 20°C. Tissue samples were sectioned at 14 μm and frozen at −20°C after drying at RT for at least one hour. Antigen retrieval was performed by immersing the sections in 1× Target Retrieval Solution (Dako, S1699) in water for 20 min, pre-heated at 80°C. Sections were washed three times in PBS containing 0.1% Tween-20 (PBSt), incubated at 4°C overnight with primary antibodies diluted in PBSt and coverslipped with parafilm. Finally, sections were washed in PBSt and incubated with secondary antibodies diluted in PBSt at RT for one hour, washed again three times in PBSt and mounted using Fluorescent mounting medium (Dako, #S3023).

#### Primary antibodies

Goat anti-GFP (1:500, Abcam, #ab6662), mouse anti-Neurofilaments (1:100, clone 2H3, DSHB), goat anti-SOX10 (1:500, Santa-Cruz, #sc-17342), mouse anti-SOX10 (1:500, Santa-Cruz, #sc-374170), rabbit anti-TH (1:1000, Pel-Freez Biologicals, #P40101-150).

DAPI (Thermo Fisher Scientific, 1:10,000, #D1306) was diluted in PBS and applied on sections for 20 min at 20-25 °C, after immunohistochemistry.

For detection of the primary antibodies, secondary antibodies raised in donkey and conjugated with Alexa-488, −555 and −647 fluorophores were used (1:1000, Molecular Probes, ThermoFisher Scientific).

#### Microscopy

Images were acquired using LSM 710 and LSM 780 Zeiss confocal microscopes equipped with 20×, 40× and 63× objectives. Images were acquired in the .lsm format and processed with ImageJ or IMARIS (8.0).

### Dataset information

Circadian cycle (bulk RNA-seq). The bulk RNA-seq data on the mouse liver circadian variation was taken from (*21*). The estimation was based on the exonic/intronic read annotation as described in the original publication.

Chromaffin differentiation (SMART-seq2). The SMART-seq2 data on the chromaffin cell differentiation was taken from (*22*).

Mouse bone marrow (in Drop). The data on the mouse bone marrow dataset is described in (*25*).

Mouse P0 and P5 hippocampus (10X Chromium) described in Hochgerner et al. (Nature Neuroscience, in review). We used P0 and P5 timepoints from Dataset C of that paper. The data is available for reviewers at GEO accession GSE104323 using token “utgraosyztctbcp”

Human postconception week 10 forebrain (10X Chromium), will be made available at GEO.

### Code availability

The software described in this paper, in the form of a pipeline called Velocyto (velox, quick; κύτoς, cell) is available at http://velocyto.org. This includes complete analysis libraries in R and Python, as well as R and Python notebooks.

## Author Contributions

S.L. conceived of the concept of RNA velocity and P.V.K. showed that RNA velocity could be detected through analysis of unspliced transcripts in single cells. P.V.K. and S.L. designed and supervised the study. P.V.K., S.L. and G.L.M. developed the analytical framework, analyzed data, made figures and drafted the manuscript, with contributions from all coauthors. P.V.K., G.L.M., R.S. and P.L. implemented the software pipeline, with assistance from V.P. and J.F. Z.L. examined RNA degradation signals. A.Z. and H.H. performed the mouse hippocampus experiment. M.E.K. and I.A. have performed and interpreted validations of chromaffin differentiation rate. E.S. provided human embryonic brain tissue. D.B. performed the human forebrain single-cell RNA-seq experiment, under supervision of G.C.B. J.G. assisted with measurement and interpretation of mouse bone marrow. The paper was read and approved by all co-authors.

## References

1. S. Linnarsson, S. A. Teichmann, Single-cell genomics: coming of age. Genome Biol 17, 97 (2016). doi: 10.1186/s13059-016-0960-x.

2. M. B. Woodworth, K. M. Girskis, C. A. Walsh, Building a lineage from single cells: genetic techniques for cell lineage tracking. Nat. Rev. Genet. 18, 230–244 (2017). doi: 10.1038/nrg.2016.159.

3. S. Skylaki, O. Hilsenbeck, T. Schroeder, Challenges in long-term imaging and quantification of single-cell dynamics. Nat. Biotechnol. 34, 1137–1144 (2016). doi: 10.1038/nbt.3713.

4. Y. S. Ju et al., Somatic mutations reveal asymmetric cellular dynamics in the early human embryo. Nature 543, 714–718 (2017). doi: 10.1038/nature21703.

5. M. A. Lodato et al., Somatic mutation in single human neurons tracks developmental and transcriptional history. Science 350, 94–98 (2015). doi: 10.1126/science.aab1785.

6. L. Haghverdi, M. Büttner, F. A. Wolf, F. Buettner, F. J. Theis, Diffusion pseudotime robustly reconstructs lineage branching. Nat. Methods 13, 845–848 (2016). doi: 10.1038/nmeth.3971.

7. M. Setty et al., Wishbone identifies bifurcating developmental trajectories from single-cell data. Nat. Biotechnol. 34, 637–645 (2016). doi: 10.1038/nbt.3569.

8. C. Trapnell et al., The dynamics and regulators of cell fate decisions are revealed by pseudotemporal ordering of single cells. Nat. Biotechnol. 32, 381–386 (2014). doi: 10.1038/nbt.2859.

9. R. Cannoodt, W. Saelens, Y. Saeys, Computational methods for trajectory inference from single-cell transcriptomics. Eur J Immunol 46, 2496–2506 (2016). doi: 10.1002/eji.201646347.

10. S. C. Bendall et al., Single-cell trajectory detection uncovers progression and regulatory coordination in human B cell development. Cell 157, 714–725 (2014). doi: 10.1016/j.cell.2014.04.005.

11. J. C. Burns, M. C. Kelly, M. Hoa, R. J. Morell, M. W. Kelley, Single-cell RNA-Seq resolves cellular complexity in sensory organs from the neonatal inner ear. Nat Commun 6, 8557 (2015). doi: 10.1038/ncomms9557.

12. J. T. Gaublomme et al., Single-Cell Genomics Unveils Critical Regulators of Th17 Cell Pathogenicity. Cell 163, 1400–1412 (2015). doi: 10.1016/j.cell.2015.11.009.

13. C. Weinreb, S. Wolock, B. K. Tusi, M. Socolovsky, A. M. Klein, Fundamental limits on dynamic inference from single cell snapshots. bioRxiv, (2017). doi: 10.1101/170118.

14. G. Schiebinger et al., Reconstruction of developmental landscapes by optimal-transport analysis of single-cell gene expression sheds light on cellular reprogramming. bioRxiv, (2017). doi: 10.1101/191056.

15. A. Zeisel et al., Coupled pre-mRNA and mRNA dynamics unveil operational strategies underlying transcriptional responses to stimuli. Mol. Syst. Biol. 7, 529 (2011). doi: 10.1038/msb.2011.62.

16. S. Picelli et al., Smart-seq2 for sensitive full-length transcriptome profiling in single cells. Nat. Methods 10, 1096–1098 (2013). doi: 10.1038/nmeth.2639.

17. S. Islam et al., Quantitative single-cell RNA-seq with unique molecular identifiers. Nat. Methods 11, 163–166 (2014). doi: 10.1038/nmeth.2772.

18. A. M. Klein et al., Droplet barcoding for single-cell transcriptomics applied to embryonic stem cells. Cell 161, 1187–1201 (2015). doi: 10.1016/j.cell.2015.04.044.

19. G. X. Y. Zheng. et al., Massively parallel digital transcriptional profiling of single cells. Nat. Commun. 8, 14049 (2017). doi: 10.1038/ncomms14049.

20. D. Gaidatzis, L. Burger, M. Florescu, M. B. Stadler, Analysis of intronic and exonic reads in RNA-seq data characterizes transcriptional and post-transcriptional regulation. Nat. Biotechnol. 33, 722–729 (2015). doi: 10.1038/nbt.3269.

21. C. Vollmers et al., Circadian oscillations of protein-coding and regulatory RNAs in a highly dynamic mammalian liver epigenome. Cell Metab. 16, 833–845 (2012). doi: 10.1016/j.cmet.2012.11.004.

22. A. Furlan et al., Multipotent peripheral glial cells generate neuroendocrine cells of the adrenal medulla. Science 357, (2017). doi: 10.1126/science.aal3753.

23. E. Gebara et al., Heterogeneity of Radial Glia-Like Cells in the Adult Hippocampus. Stem Cells 34, 997–1010 (2016). doi: 10.1002/stem.2266.

24. R. J. Johnston, Jr, C. Desplan, Stochastic Mechanisms of Cell Fate Specification that Yield Random or Robust Outcomes. Annu. Rev. Cell Dev. Biol. 26, 689 (2010). doi: 10.1146/annurev-cellbio-100109-104113.

25. V. Petukhov et al., Accurate estimation of molecular counts in droplet-based single-cell RNA-seq experiments. bioRxiv, (2017). doi: 10.1101/171496.

